# E(spl)m4 Directly Antagonizes Traf4 to Inhibit JNK Signaling in *Drosophila*

**DOI:** 10.1101/2025.09.29.679213

**Authors:** Katrin Strobel, Jennifer Falconi, Cédric Leyrat, Rémi Logeay, Sarah J. Bray, Alexandre Djiane

## Abstract

TRAF proteins are adaptor proteins that participate in signal transduction downstream of the Toll or TNF receptors and could elicit E3-Ubiquitin Ligase activity. They have been implicated in multiple processes during signal transduction, inflammation, and morphogenesis. In Drosophila, Traf4 has been implicated in the regulation of JNK signaling and cell death as well as Adherens Junctions regulation. Using overexpression approaches, we show here that Traf4 promotes JNK and caspase activation, as well as junctional E-Cadherin/ β-Catenin depletion, resulting in epithelial cell delamination. Using biochemical, modelling, and functional genetics approaches, we further show that the Bearded-type small protein E(spl)m4 binds to Traf4, and inhibits its downstream signaling towards JNK activation and cell delamination, without affecting the effects of Traf4 on Adherens Junctions. Thus, this study identifies an endogenous peptide inhibiting Traf4 signaling, potentially by blocking Traf4 trimerization.

## INTRODUCTION

TRAF proteins form a family of adaptor proteins implicated in the transduction of signaling pathways. Initially identified associated with the TNF Receptor (hence their name of TNF Receptor Associated Factor), they can associate with various receptors including Toll receptors, and mainly activate the JNK and NF-kB pathway (Bradley & Pober, 2001; Chung *et al*, 2002). They have been implicated at multiple steps during inflammation and immunity (Dhillon *et al*, 2019; Lalani *et al*, 2018). TRAFs all share a common C-terminal domain, the MATH/TRAF domain, important for protein-protein interaction with surface receptors (e.g. TNFR), specific adapters (e.g. TRADD) or downstream effectors. The MATH domain also mediates the trimerization of TRAF proteins (Bradley & Pober, 2001; Chung *et al*, 2002; Pullen *et al*, 1999; Kim *et al*, 2016; Baud *et al*, 1999; Park, 2018). TRAFs can also contain N-terminal RING and Zn fingers which are important for their function and could thus elicit E3-Ubiquitin Ligase activity (e.g. TRAF2 and TRAF6) (Middleton *et al*, 2017).

In *Drosophila* there are three TRAFs named after their closest mammalian orthologue: Traf-like, Traf4 (formerly known as dTraf1), and Traf6 (formerly known as dTraf2) (Grech *et al*, 2000). Traf4 contains several zinc fingers including some RING-type zinc fingers and a MATH/TRAF domains (Fig. 1A). The presence of RING fingers suggests a potential E3-Ubiquitin Ligase activity, but this has not been formally tested. Even though Traf4 can bind to Pelle and activate NF-kB pathways (Zapata *et al*, 2000), it has been mainly implicated in the regulation of JNK signaling and in the promotion of apoptosis (Cha *et al*, 2003; Kuranaga *et al*, 2002; Liu *et al*, 1999; Lu *et al*, 2017). The Traf4-mediated JNK activation occurs via its interaction with the upstream MAPKKK Misshapen, Ask1, and Tak1 (Geuking *et al*, 2005; Liu *et al*, 1999; Kuranaga *et al*, 2002), and the consequences to this JNK activation are context dependent (Agrawal *et al*, 2016; Kuranaga *et al*, 2002; Lu *et al*, 2017). Traf4 has also been shown to regulate morphogenesis. Indeed, Traf4 binds to and appears to destabilize Armadillo (the *Drosophila* homologue of β-Catenin), and controls Adherens Junctions polarization and dynamics during embryonic mesoderm involution (Mathew *et al*, 2009, 2011). This junction dynamics activity of Traf4 appears conserved during evolution and the human TRAF4 has been shown to control apical Tight Junctions stability and cell migration in cultured cell lines (Rousseau *et al*, 2013; Wang *et al*, 2013). Traf4 also binds to Bazooka and controls asymmetric cell division in *Drosophila* embryonic neuroblasts (Wang *et al*, 2006). Whether the role of Traf4 on membrane determinants dynamics (Arm, Baz…) is linked to its effect on JNK signaling, or whether these activities are independent remains unexplored.

**Figure 1.**
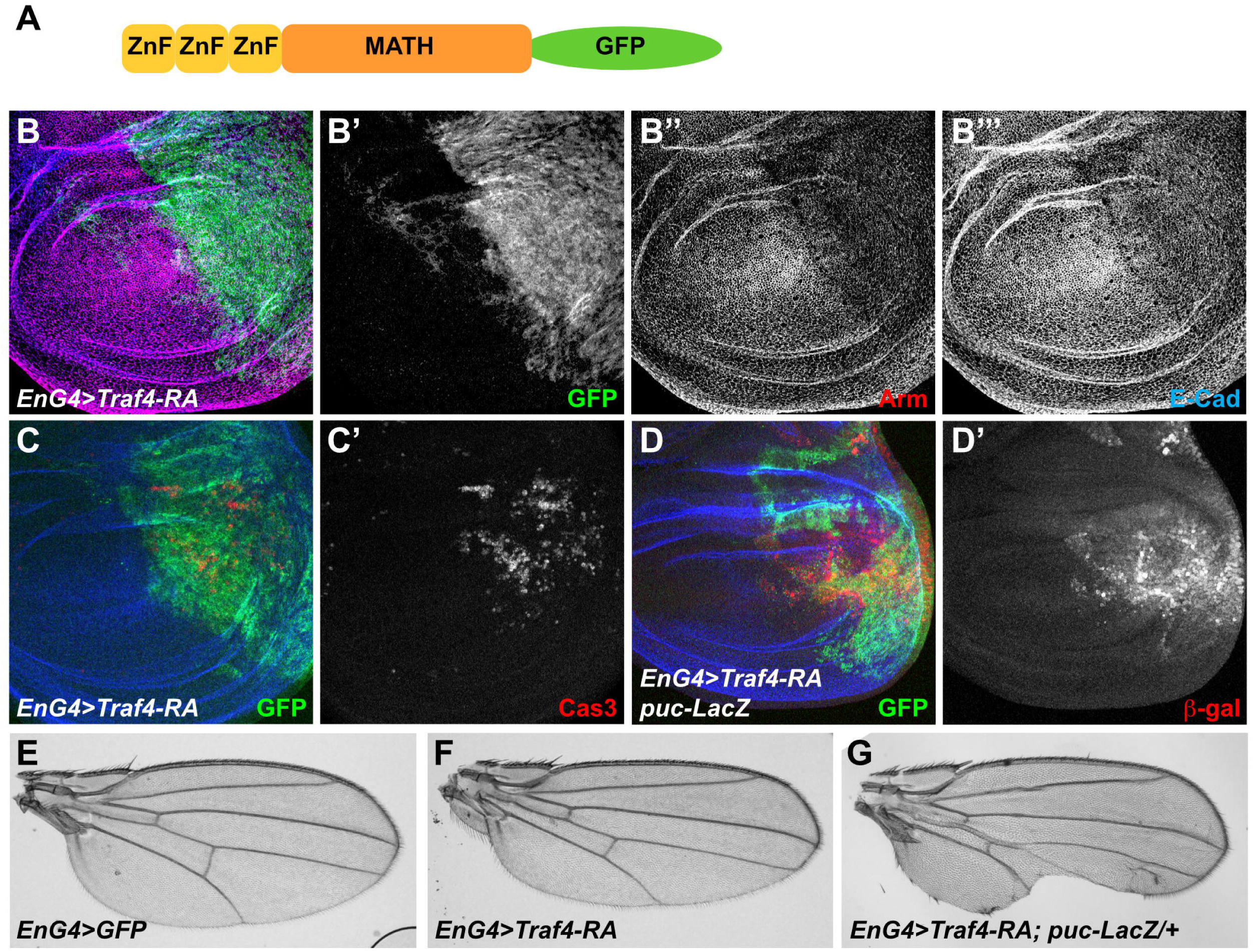
Overexpressed Traf4 affects Adherens Junctions and promotes JNK activity. **A.** Schematic of Traf4-PA structure with N-terminal zinc finger domains (ZnF - yellow boxes), and C-terminal MATH domain (orange box). The C-terminal GFP tag is represented in green. **B-C.** Staining of 3^rd^ instar wing imaginal disc with overexpressed Traf4-PA:GFP in the posterior compartment showing the Traf4-PA:GFP overexpression domain (GFP, green in B&C, white in B’), the Adherens Junctions components Armadillo (Arm, red in B, white in B’’) and E-Cadherin (E-Cad, blue in B&C, white in B’’’), and cleaved Caspase 3 (Cas3, red in C, white in C’). **D.** Staining of 3^rd^ instar wing imaginal disc in the *puc-LacZ/+* genetic background with overexpressed Traf4-PA:GFP in the posterior compartment showing Traf4-PA:GFP (GFP, green in D), E-Cadherin (blue in D), and JNK activation (β-gal, red in D, white in D’). **E-G.** Female adult wings of the indicated genotypes showing the smaller posterior compartment after Traf4-RA overexpression (E&F) and the loss of posterior wing disc material in the *puc-LacZ/+* background (G).

The activity of signaling pathways transduction cascades is tightly regulated ensuring fine tuning and preventing their inappropriate firing. This is classically achieved through intricate post-translational modifications, in particular as part of feed-back loops or cross-talks between different pathways. Additionally, signaling pathways can be regulated by protein/protein interactions. In the case of the adaptor proteins Trafs, in mammals they interact with the upstream receptors either directly (TNFR2, CD40) or indirectly via TRADD (TNFR1) or MyD88-IRAK (Toll-Like Receptors), and bring in the complex downstream effectors of the JNK and NF-kB signaling pathways, MAP3Ks or IKK respectively (reviewed in (Bradley & Pober, 2001; Chung *et al*, 2002; Park, 2018). The trimerization of Trafs through their MATH domains has been shown as a key step for the downstream NF-kB and JNK pathway activation (Baud *et al*, 1999). But, protein interactions can also be inhibitory, for instance by preventing signaling complex formation, or by preventing the interaction between an enzyme and its substrate during regulatory post-translational modifications.

Recent studies have highlighted the existence of hundreds of small open-reading frames and peptides (<100 aa) and seminal studies suggest that they could act as potent protein activity regulators. For instance Pri peptides have been shown to interact with the E3-Ubiquitin Ligase Ubr3 to regulate its activity and epidermal patterning (Markus *et al*, 2023). Similarly, Sarcolamban control the activity of the Ca2+ transporter SERCA and thus different aspects of calcium signaling (Magny *et al*, 2013). Even though slightly above the 100 aa cut off, the Bearded proteins (circa 150 aa) also play important regulatory roles, in particular during Notch signaling. Bearded proteins are encoded by the bearded genes, a family of closely related paralogues grouped in two genomic clusters. The first cluster contains the genes *BobA*, *Tom*, *Brd*, and *Ocho*; the second cluster contains the genes *E(spl)mα, m2, m4*, and *m6* which are interspersed with the more classical Notch target genes coding for the transcription repressors of the E(spl)-HLH family. All Bearded proteins except E(spl)m2 interact with the E3-Ubiquitin Ligase Neuralized (Neur), and Tom has been shown to inhibit Neur activity resulting in decreased ubiquitylation of the Notch ligand Delta and thus lower Notch pathway activity (Bardin & Schweisguth, 2006). Since the interaction between Bearded proteins and Neur occurs at the level of the highly conserved 10aa long motif 2 (Bardin & Schweisguth, 2006), it is assumed that Bearded proteins act similarly to Tom. Indeed, gain of function mutations in the *Brd* gene lead to classical Notch inhibition phenotypes with supernumerary sensory bristles, resembling *neur* or *Dl* loss of function mutations. This redundancy between Bearded proteins might explain why loss of function mutants have no apparent phenotypes (Chanet *et al*, 2009). Importantly, the antagonistic interaction between Bearded proteins and Neur is also found in other contexts besides Notch signaling regulation, such as during morphogenetic movements and involution of the embryonic mesoderm (Chanet & Schweisguth, 2012; Perez-Mockus *et al*, 2017).

Studying the function of Traf4 during Adherens Junctions (AJ) regulation and JNK pathway activation in *Drosophila* wing imaginal discs, we identify here the small protein E(spl)m4 as a binding inhibitor of Traf4 activity. We further show that E(spl)m4 only inhibits a subset of Traf4 activities, namely its role in JNK signaling and caspase activation, but appear inefficient on Traf4-mediated AJ material destabilization. Finally we provide protein structure modelling suggesting the inhibitory role of E(spl)m4 is achieved by preventing Traf4 trimerization.

## RESULTS

### Traf4 overexpression destabilizes AJ

*Traf4* encodes for an adapter protein belonging to the family of the TNF receptor associated factors, and Traf4-PA, the main Traf4 isoform contains a N-terminal unstructured domain (circa 125 aa), three 3 zinc finger domains (related to the RING/FYVE/PHD zinc finger domains), and a large C-terminal MATH/TRAF domain implicated in the interaction with the TNF receptors (Fig. 1A) (Bradley & Pober, 2001; Chung *et al*, 2002; Grech *et al*, 2000). Traf4 is detected at sites of cell-cell adhesion, including the tight junctions in mammalian epithelia (Rousseau *et al*, 2013; Wang *et al*, 2013), and the Adherens Junctions (AJ) in *Drosophila* embryos. Loss of *Traf4* in the latter case results in elevated accumulation of the AJ component Armadillo / β-Catenin and altered morphogenetic movements (Mathew *et al*, 2009, 2011). In order to investigate whether Traf4 has a more widespread role in regulating AJ associated proteins, in particular the Cadherin/Catenin complex, we overexpressed a C-terminally GFP tagged Traf4 in the wing imaginal disc epithelial cells, using the *engrailed-Gal4* driver (*en-Gal4*). Traf4 overexpression led to a dramatic reduction of the AJ associated components E-Cadherin (E-Cad) and its partner Armadillo (Arm; β-Catenin in vertebrates; Fig. 1B). We observed extremely discreet effects on aPKC (slight decrease) present at the apical and sub-apical membrane regions and on the Septate Junction associated protein Discs Large (Dlg; slight increase) after Traf4 overexpression, possibly reflecting indirect effects resulting from the balance between the different lateral membrane compartments (Supplemental Fig. S1). These results show that Traf4 overexpression in the *Drosophila* wing disc epithelium leads to defects in the apical compartment of epithelial cells, and in particular the AJ, a role consistent with its described role on AJ regulation during mesoderm invagination during early embryogenesis (Mathew *et al*, 2009, 2011).

Traf scaffolds have been implicated in signal transduction downstream of TNF receptors, typically mediating NF-kB and JNK activation (Bradley & Pober, 2001; Chung *et al*, 2002). In *Drosophila*, Traf4 has been shown to activate robustly JNK and to promote cell death (Cha *et al*, 2003; Kuranaga *et al*, 2002; Liu *et al*, 1999; Lu *et al*, 2017). To reproduce these results, we thus examined the expression of the JNK activity reporter *puckered-LacZ (puc-LacZ)*. Overexpression of Traf4, triggered in a strong increase in *puc-LacZ* expression (Fig. 1D) which was only detected within the Traf4-overexpressing cells, suggesting that the effects of Traf4 on JNK signaling were cell autonomous. A large portion of these *puc-LacZ* positive Traf4-overexpressing cells delaminated and were found basal to the main epithelial sheet. This correlated with misshapen adult wings, with the posterior *en-Gal4* domain reduced in size, dented and lacking portions of the wing margin (Fig. 1E&G). It should be noted that the *puc-LacZ* reporter represents a sensitized JNK background and the sole overexpression of *Traf4* under the *en-Gal4* driver resulted in a much milder phenotype even though increased caspase could be seen (Fig. 1C) with correct overall wing shape and posterior margin, but with reduced posterior compartment size (Fig. 1E&F).

### Traf4 overexpression promotes epithelial delamination

The genetic interaction between *Traf4* and *puc-LacZ* suggested that Traf4 over-expression could promote epithelial delamination. To test this, we overexpressed Traf4 at high levels and in a more restricted number of cells using the *patched-Gal4* driver (*ptc-Gal4*) along the antero-posterior boundary of wing imaginal discs at 29°C. Under these conditions, Traf4 overexpressing cells delaminated from the epithelium, and were found basally (Fig. 2). These delaminated cells also migrated towards the posterior part of the disc and expressed activated cleaved caspase 3. The activation of the effector caspases was at least in part required for the delamination and migration of Traf4 overexpressing cells, as co-expressing the effector caspases inhibitor P35, which blocks the cascade downstream of caspase 3/drICE, suppressed the delamination phenotype (even though cells were still positive for the active cleaved caspase 3, showing diffuse cytoplasmic staining; Fig. 3A-D).

**Figure 2.**
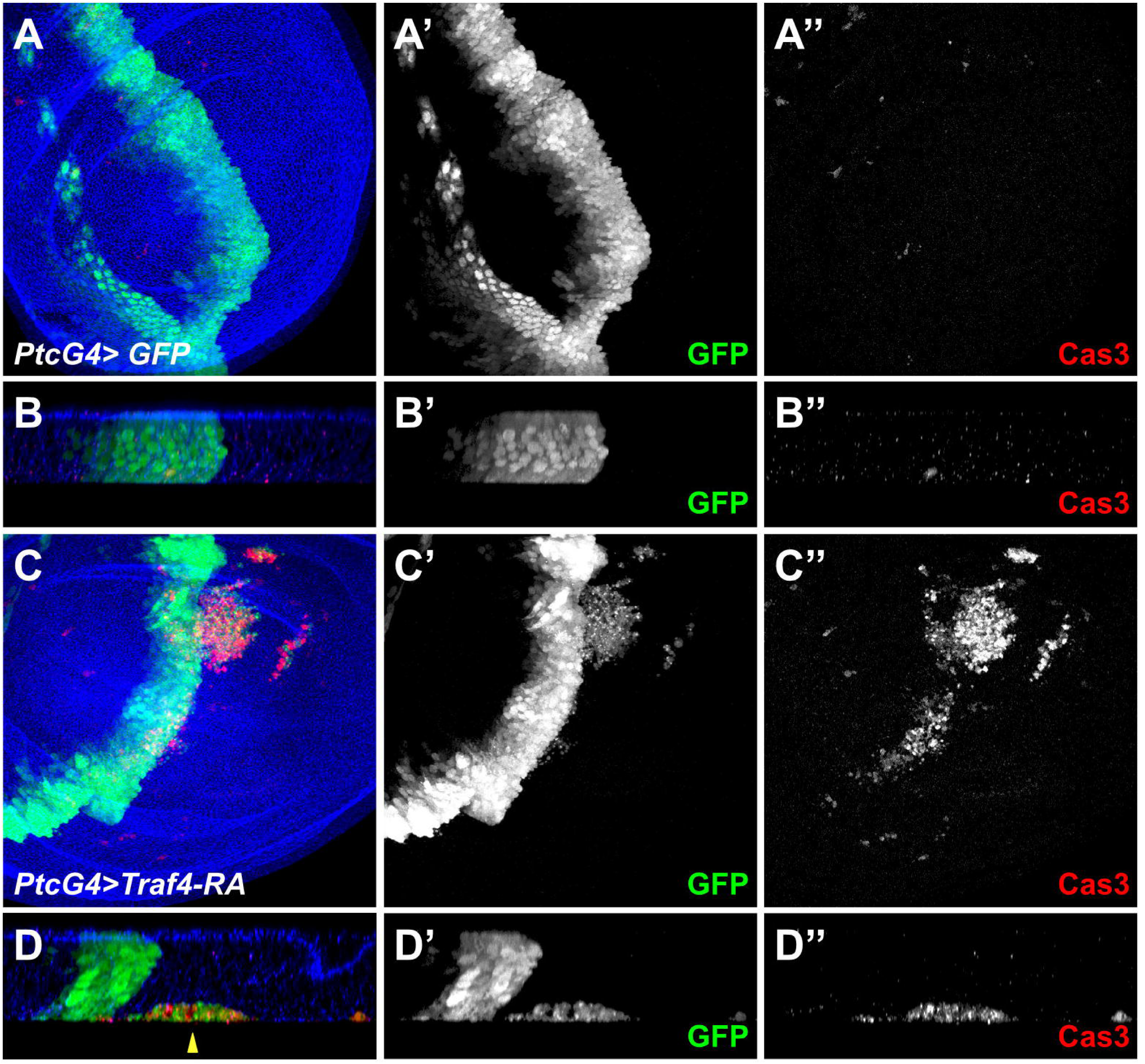
Overexpressed Traf4 promotes cell delamination. **A-D.** Staining of 3^rd^ instar wing imaginal disc overexpressing either GFP (A&B) or Traf4-RA:GFP (C&D) and showing the overexpression domain (GFP, green in A-D, white in A’-D’), E-Cadherin (blue in A-D), and cleaved Caspase 3 (Cas3, red in A-D, white in A’’-D’’). Discs are shown along the xy axis in A&C and along the z axis in B&D. Note the accumulation of GFP+/Cas3+ cells on the basal side of the discs (yellow arrow, opposite to E-Cadherin) after Traf4-RA overexpression at 29°C.

**Figure 3.**
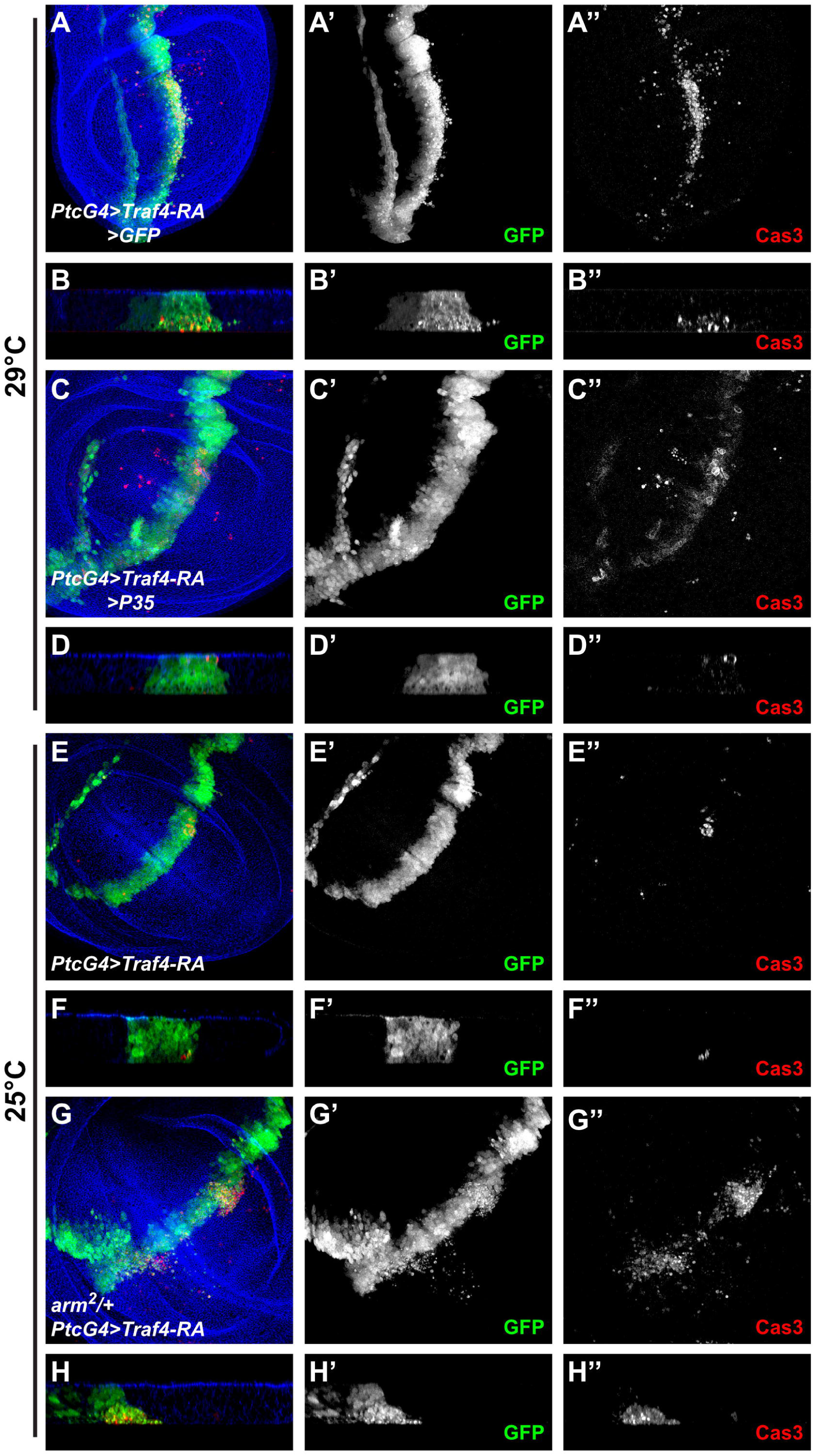
Traf4-induced cell delamination requires caspase activity. **A-D.** Staining of 3^rd^ instar wing imaginal disc overexpressing Traf4-RA:GFP at 29°C either with GFP (A&B) or with the caspase inhibitor P35 (C&D) and showing the overexpression domain (GFP, green in A-D, white in A’-D’), E-Cadherin (blue in A-D), and cleaved Caspase 3 (Cas3, red in A-D, white in A’’-D’’). Discs are shown along the xy axis in A&C and along the z axis in B&D. **E-H.** Staining of 3^rd^ instar wing imaginal disc overexpressing Traf4-RA:GFP at 25°C either in wild type background (E&F) or with one mutant copy for armadillo (*arm^2^/+*; G&H) and showing the overexpression domain (GFP, green in E-H, white in E’-H’), E-Cadherin (blue in E-H), and cleaved Caspase 3 (Cas3, red in E-H, white in E’’-H’’). Discs are shown along the xy axis in E&G and along the z axis in F&H.

The down-regulation of AJ was also critical for the delamination of Traf4 overexpressing cells. When overexpressed with *ptc-Gal4* but at lower levels (shifting the flies at 25°C instead of 29°C), Traf4 was unable to induce cell delamination. However, cell delamination could be observed under these conditions in animals with half the normal dosage for *armadillo* (*arm* heterozygous mutants; Fig. 3E-H).

Altogether these results suggest that cells overexpressing Traf4 have weakened AJs and are prone to epithelial delamination and subsequent basal migration. In the process, these cells activate the JNK pathway and the pro-apoptotic caspase cascade. It is noteworthy here, that blocking apoptosis by P35 suppresses Traf4-mediated delamination.

### The bearded-type peptide E(spl)m4 binds to Traf4

We then searched for partners of Traf4 which could be important for its function. Mining the genome-wide protein-protein interaction databases highlighted the E(spl)m4 as a high confidence Traf4 interactor (Giot *et al*, 2003; Tang *et al*, 2023). *E(spl)m4* is part of a family of genes encoding short (circa 150aa) polypeptides: the Bearded proteins (Supplemental Fig. S2A). The eight Bearded genes are divided in two clusters: *E(spl)mα, m2, m4*, and *m6*, are found in the Enhancer of split (E(spl)) complex intertwined with the canonical Notch targets encoding bHLH transcriptional repressors on the 3R chromosome; the other four, *BobA, Tom, Brd*, and *Ocho,* are found in a separate cluster on the 3L chromosome. They have all been shown to interact with the E3-Ubiquitin Ligase Neuralized and at least Tom has been shown to inhibit Neuralized activity during Notch signaling by competing with Dl to bind to Neuralized (Bardin & Schweisguth, 2006).

We therefore tested whether the Bearded family proteins, and more specifically E(spl)m4 could bind to Traf4. Using yeast two-hybrid (Y2H), we detected a strong interaction between E(spl)m4 and Traf4-RA (Fig. 4A), thus validating the previously identified interaction. A slightly weaker interaction was also detected with E(spl)m2. We could not detect any interaction by Y2H for any of the other Bearded peptides, even for the E(spl)m4 very close paralogue Tom (Fig. 4A).

**Figure 4.**
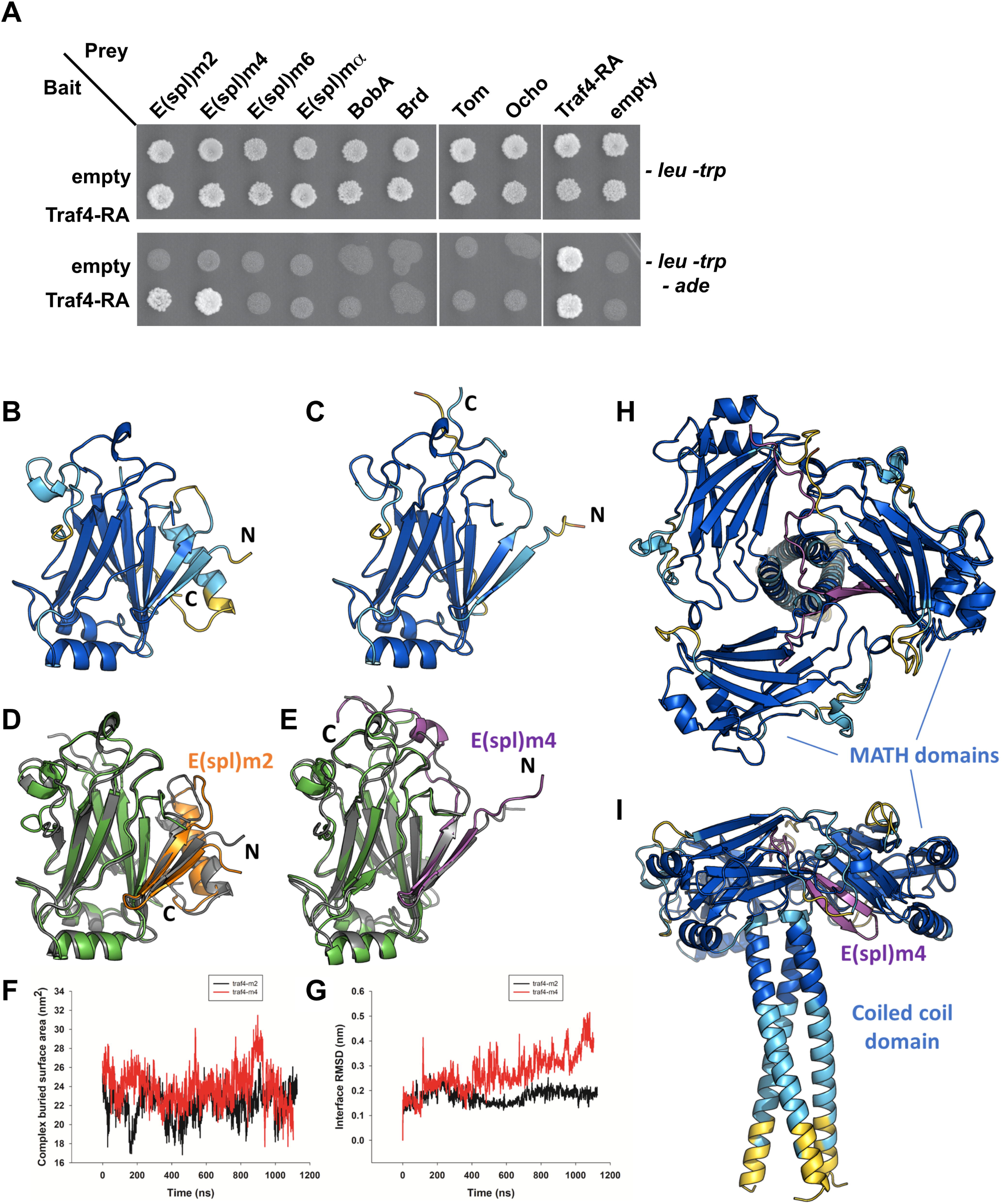
E(spl)m4 and E(spl)m2 small proteins interact with Traf4. **A.** Yeast two-hybrid assay using Traf4-RA as a bait and the different Bearded proteins as preys. Upper panel: selection of dicaryonic yeast cells containing both bait and prey vectors on media lacking leucine and tryptophan *(-leu -trp*). Lower panel: interaction assay through adenosine auxotrophy rescue by growing the dicaryons on media lacking leucin, tryptophan, and adenosine *(- leu -trp -ade*). Empty vectors are used as negative controls testing for potential self-activation of either the baits or the preys. Here the Traf4-RA prey vector is self-activating. **B-I.** Predicted structures and molecular dynamics simulations (MDS) of Traf4-E(spi)-m2 and Traf4-E(spi)-m4 complexes **B-C.** Alphafold3 (AF3) structural model of Traf4-E(spl)-m2 (B) and Traf4-E(spl)-m4 (C), colored using the standard AF3 confidence (pLDDT) color scale (blue = very high, cyan = confident, yellow = low). **D-E.** Superimposition of Traf4-E(spl)-m2 (D) and Traf4-E(spl)-m4 (E) structural models at the beginning and at t=1µs of the MDS trajectory. The starting model is colored in dark grey, while the t=1µs snapshot is colored by chain (green for Traf4 MATH domain and orange and magenta for E(spl)-m2 and E(spl)-m4, respectively). **F.** Interface buried surface area of the Traf4-E(spl)-m2 (black curve) and Traf4-E(spl)-m4 complexes (red curve) as a function of simulation time. **G.** Interface root mean square deviation (iRMSD) of the Traf4-E(spl)-m2 (black curve) and Traf4-E(spl)-m4 complexes (red curve) as a function of simulation time. **H-I.** Top (H) and side (I) views of an AF3-predicted structural model of trimeric Traf4 superimposed with the predicted Traf4-E(spl)-m4 complex, highlighting the steric incompatibility of the two interactions. The Traf4 trimer and E(spl)-m4-bound MATH domain are colored using the standard AF3 color scale, and E(spl)-m4 is colored in magenta.

To investigate a potential direct interaction between Traf4 and E(spl)m2 or E(spl)m4, we used Alphafold3 protein structure prediction, which yielded high confidence models for complexes consisting of Traf4 MATH domain (residues 328-480) and regions of E(spl)m2 and E(spl)m4 encompassing residues 175-216 and residues 100-136, respectively (Fig. 4B&C). The core interacting motif corresponds to a β-hairpin containing a conserved GTFFW sequence that packs against the first β-strand of Traf4 MATH domain. This motif corresponds to the previously highlighted motif of homology 3 amongst Brd proteins and is different from the motif 2 mediating the interaction with Neuralized (Supplemental Fig. S2A). The average pLDDT values across the core interface residues of the predicted complexes are respectively 81.4 and 86.8 for E(spl)m2 and E(spl)m4, suggesting stable complexes. Despite the core GTFFW motif being conserved in several other Bearded proteins (e.g. E(spl)mα, Tom, Ocho), there is significant divergence in the N- and C-flanking sequences (Supplemental Fig. S2A). Accordingly, predictions of Traf4 in complex with E(spl)mα, Tom, and Ocho resulted in models with poor confidence scores and different binding poses, in agreement with the lack of interaction observed in Y2H experiments.

In order to confirm the stability of the Traf4-E(spl)m2 and -E(spl)m4 complexes, we turned to classical molecular dynamics simulations (Fig. 4D-G). Both complexes were stable during the simulations, showing only minor fluctuations of the interface buried surface area (BSA) and interface RMSD (iRMSD) over time (Fig. 4F&G). Somewhat higher deviations of iRMSD were observed for Traf4-E(spl)m4, which originated from conformational rearrangements of residues 121-134 of E(spl)m4 (Fig. 4G).

Traf proteins have been shown to form trimers at the level of their MATH domain, which is required for the downstream activation of the NF-kB and JNK pathways following Receptor/Traf complex formation (Bradley & Pober, 2001; Chung *et al*, 2002; Pullen *et al*, 1999; Kim *et al*, 2016; Baud *et al*, 1999; Park, 2018). Interestingly, the interaction between Traf4 and E(spl)m4 (and E(spl)m2) occurs at the level of the MATH box and is predicted to disrupt/prevent Traf4 trimerization (Fig. 4H&I), suggesting that E(spl)m4 could represent a Traf4 inhibitor. More specifically, first β-strand of the MATH domain (residues 332-338) was predicted to be engaged with E(spl)m4 (Fig. 4C). Comparison of the Traf4 - E(spl)m4/ E(spl)m2 models with available crystal structures of mammalian TRAFs complexes indicates that the interface on Traf4 is distinct from interfaces involved in interactions with the upstream activating receptors or adaptors (e.g. TNFR2, TRADD; Fig. S2B).

### The bearded-type peptide E(spl)m4 inhibits Traf4 effect on epithelial delamination

We thus asked whether the Traf4/ E(spl)m4 interaction is relevant functionally. Thus, we studied the effect of co-expressing under the *en-Gal4* driver, E(spl)m4 or other Bearded family peptides together with Traf4-PA. We investigated whether E(spl)m4 was able to inhibit the effect of Traf4-PA overexpression on E-Cad levels and JNK pathway activation in the 3rd instar larval wing discs. When co-expressed with Traf4-PA in the background of the *puc-LacZ* JNK activity reporter line at 25C, E(spl)m4 was able to inhibit *puc-LacZ* expression (in 5/8 discs compared to 0/8 with expression of a GFP control), but was completely unable to suppress the Traf4-mediated E-Cad downregulation (Fig. 5A&B). Consistently, in adult wing discs, overexpressed E(spl)m4 was able to prevent the loss of posterior tissue, while Tom was not (Fig. 5D-E). These results suggest that E(spl)m4 inhibits only a subset of the effects of overexpressed Traf4, namely its action on JNK pathway, but not its effects on E-Cad levels. Furthermore, overexpressed E(spl)m4 suppressed the Traf4-PA-induced cell delamination and caspase activation after overexpression at 29°C with the *ptc-Gal4* driver (in 8/10 discs compared to 0/10 for a GFP overexpression control; Supplemental Fig. S3). Taken together these results show that the physical interaction between E(spl)m4 and Traf4 is functional and leads to an inhibition of at least some of Traf4 activities, namely its effect on JNK pathway and caspase cascade activation.

**Figure 5.**
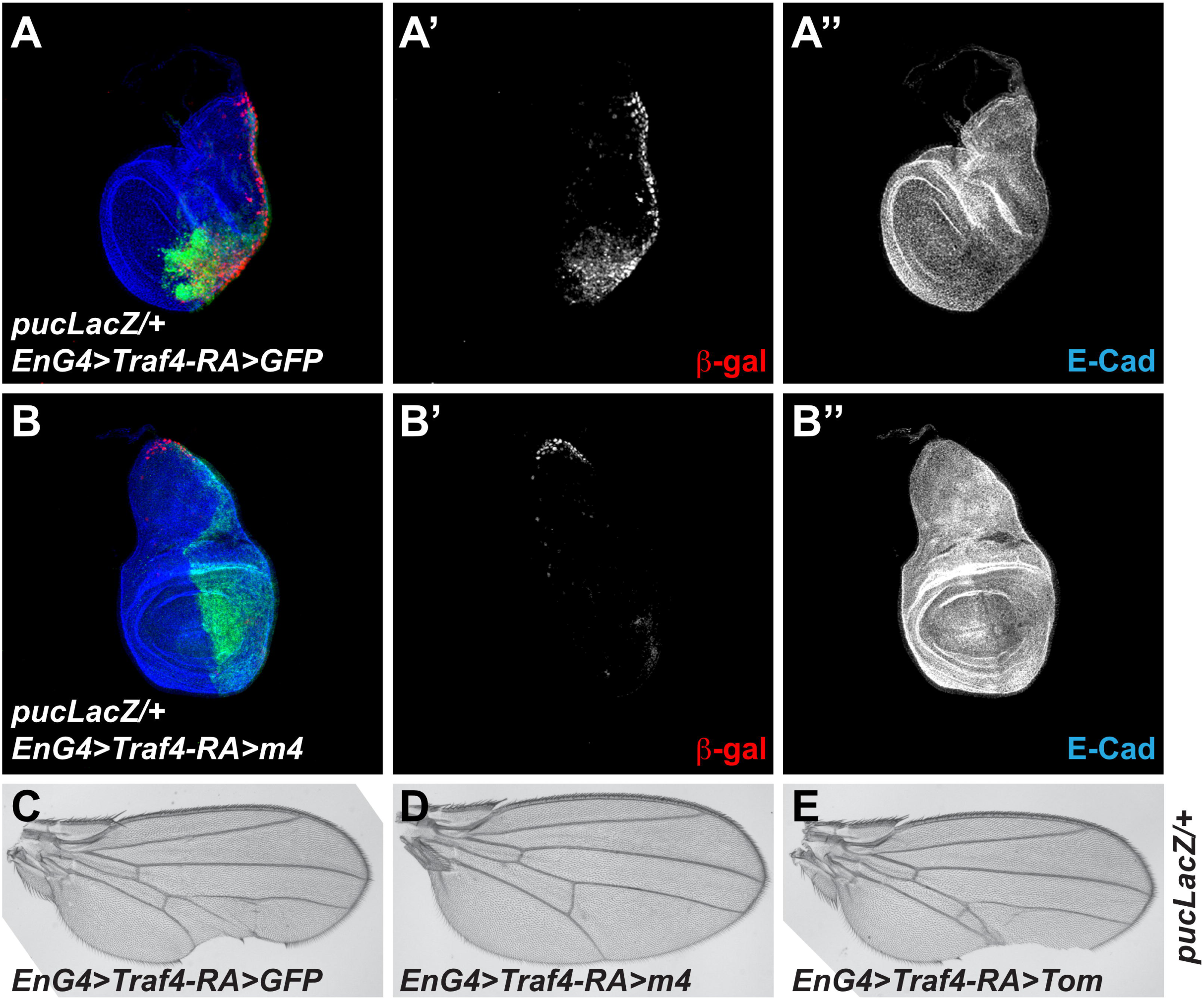
E(spl)m4 suppresses the effects of Traf4 overexpression. **>A-B.** Staining of 3^rd^ instar wing imaginal disc in the *puc-LacZ/+* genetic background and co-overexpressing Traf4-PA:GFP in the posterior compartment either with GFP (A) or E(spl)m4 (B). Discs show the Traf4-PA:GFP overexpression domain (GFP, green in A&B), JNK activation (β-gal, red in A-B, white in A’-B’), and E-Cadherin (E-Cad, blue in A-B, white in A’’-B’’). **C-E.** Female adult wings after Traf4-RA overexpression in the *puc-LacZ/+* background either with GFP (C), E(spl)m4 (D), or Tom (E).

### Overexpressed E(spl)m4 recapitulates some of Traf4 mutant defects

To further strengthen the functional antagonism between Traf4 and E(spl)m4, we thought to investigate whether overexpressed E(spl)m4 could phenocopy the loss of *Traf4*.

First, to study the function of *Traf4*, we generated new loss of function mutations. Using imprecise excision of the transposable P element *P{EP}Traf4^EP578^*, we isolated several small genomic deletions affecting the *Traf4* locus, including the *Traf4^ex111^* mutant, in which the *Traf4-RA* specific exon and the transcription and translation start sites are missing. These new mutants were likely hypomorph since they were homozygous viable. Indeed, the *Traf4* null mutant *Traf4^L2^* generated by Wang and colleagues by mobilization of *P{EP}Traf4^EP578^* removing the whole open reading frame is homozygous lethal (Fig. 6A) (Wang *et al*, 2006). *Traf4^L2^* null mutant clones died and were excluded from the wing disc epithelium, making the analysis of the mutant phenotype impossible at the cellular level. On the other hand, *Traf4^ex111^* was viable and the only visible phenotype was an improper patterning of the adult wing margin bristles (WMB), with clusters of two or three bristles being formed in all homozygous or trans-heterozygous combinations tested (Fig. 6B-D). This phenotype was also seen in the rare tiny adult *Traf4^L2^* wing clones that survived in the adult, and in *Traf4* knock-down experiments by RNAi (driven by *scabrous-Gal4; sca-Gal4*), showing that this phenotype was specific (Fig. 6E&F). These results suggest that lowering the levels of *Traf4* in the wing, leads to a problem in the specification of the number and spacing of sensory organs at the wing margin. Overexpressed E(spl)m4 in the proneural clusters using the *sca-Gal4* driver led to strong alterations of the WMB pattern, including mis-specifications with naked margin or thinner bristles, but also to clusters of WMB similar to those observed in *Traf4* mutants, albeit to a much lower penetrance (Fig. 6G). Even though, the effect of overexpressed E(spl)m4 on Delta/Notch signaling introduces confounding factors in the analysis of the WMB phenotypes, the clusters of WMB observed both in *Traf4* mutants and in overexpressed E(spl)m4, are consistent with Traf4 and E(spl)m4 exhibiting opposite roles. The lower penetrance observed with E(spl)m4 overexpression compared to Traf4 loss of function, could thus be a consequence of the interference with Notch pathway alterations. Alternatively, it could also reflects that the *Traf4* mutant WMB spacing defects result from a combined effect on both JNK signaling and Adherens Junctions (*Traf4* mutant), when E(spl)m4 overexpression is only able to prevent JNK signaling downstream of Traf4 (Fig. 6M).

**Figure 6.**
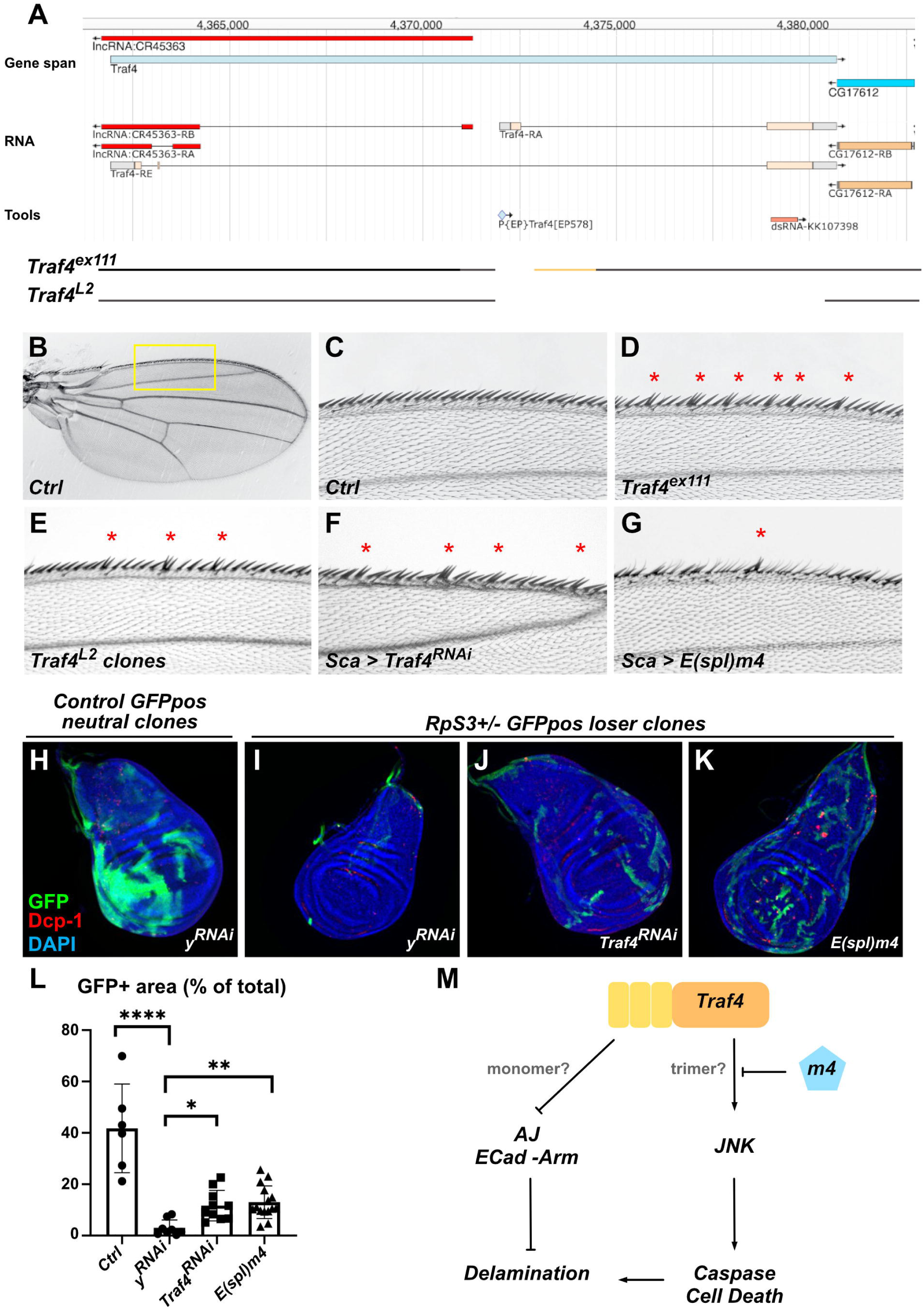
Overexpressed E(spl)m4 mimics some Traf4 loss of function phenotypes. **A.** Upper panel: Traf4 genetic locus adapted from the JBrowse from Flybase and showing the location of the P element EP578 used from generating new Traf4 alleles, and the sequences targeted by the Traf4-RNAi KK107398. Lower panel: diagram showing the extent of the Traf4^ex111^ and Traf4^L2^ deletion mutants; interrupted lines represents the lacking DNA; black lines represent the DNA still present; orange lines represent regions of uncertainty due to the PCR-based mapping. Traf4^L2^ mapping is based on published data. **B-G.** Wing margin bristles in the indicated genotypes showing defects in spacing (red stars). (B) Low magnification image of control wing. (C-G) High magnification images corresponding to the region highlighted by the yellow box in B. **H-K.** Staining of 3^rd^ instar wing imaginal disc with mosaic tissues to assay for cell competition in which clones of GFP marked clones of mutant tissues were generated and marked positively by GFP. Clones were either *wild-type* (no cell competition, H), lacking one copy of *RpS3* (*RpS3-/+* loser clones being eliminated, I), or *Rps3-/+* and overexpressing either an RNAi for *Traf4* (J) or E(spl)m4 (K). Discs show nuclei (DAPI, blue), the extent of clone growth (GFP, green), and apoptotic cells (Dcp-1, red). **L.** Quantification of the clone area from (H-K) shown as total clone area in the wing pouch normalized by total wing pouch area. n = 6-15 discs. Error-bars show standard error of the mean (sem). One-way ANOVA statistical test. **** p < 0.0001, ** p < 0.01, * p < 0.05 **M.** Model for Traf4 inhibition by E(spl)m4.

Recently, Traf4 has been implicated in the control of the loser state during Myc/Ribosomal-controlled cell competition. Cells with lower *Myc* or *Ribosomal gene* copy number are normally eliminated from the wing disc tissue when confronted with cells with higher Myc/Ribosome content. In this context, the loss of *Traf4* prevented loser cell elimination and restored the size of low *Myc* clones (Kodra *et al*, 2024). In an equivalent system, where loser cells were generated by removing one copy of *RpS3*, a gene coding for a structural protein of the small ribosomal subunit, and key dosage-sensitive gene in the context of Myc/Ribosome controlled cell competition (Kiparaki *et al*, 2022; Baumgartner *et al*, 2021), impairing *Traf4* expression prevented loser cell elimination as judged by the increase in average mutant clone size (Fig. 6H-J&L). Similarly, overexpression of E(spl)m4 prevented loser cell type elimination (Fig. 6K&L).

Together, these results thus demonstrate that overexpressed E(spl)m4 mimics the loss of function of *Traf4* and are consistent with an inhibitory role of E(spl)m4 on Traf4 in cells with endogenous levels of Traf4, extending similar conclusions between E(spl)m4 and overexpressed Traf4.

## DISCUSSION

In this study we have shown that Traf4 in *Drosophila* exhibits at least two activities in the developing wing imaginal disc: i) promoting JNK signaling and cell death, and ii) regulating AJ material abundance. We further showed that the small protein E(spl)m4 is a binding inhibitor of Traf4 preventing specifically its action on JNK signaling, but not on AJ (Fig. 6M). Based on protein interaction modelling we propose that the E(spl)m4 disrupts the trimerization of Traf4 at the level of its MATH/TRAF domain, thus inhibiting downstream JNK activation.

The interaction between Traf4 and E(spl)m4 was predicted to occur between the MATH/TRAF domain and a stretch of highly conserved residues between the different Bearded family proteins. This C-terminal motif is different from the motif 2 previously identified in Bearded proteins and mediating their interaction with Neuralized (Bardin & Schweisguth, 2006). Despite the conservation of the Traf4-interacting domain amongst Bearded proteins, we observed that the binding and the inhibition appeared restricted to E(spl)m4 (and to a lesser extent to E(spl)m2). This suggests that other residues found in other Brd proteins might destabilize the interaction. Traf4 residues interacting with E(spl)m4 are predicted to be inside the MATH/TRAF domain trimer thus potentially destabilizing the Traf4 aggregate and further pathway transduction. This binding and inhibition mechanism appear thus different from already known TRAF inhibitors which were predicted to interact on the outside of the MATH/TRAF domain trimer and masking their interaction with their upstream receptors such as the interaction between CD40 and TRAF6 inhibited by the TRAF-STOP inhibitor 6877002 (Zarzycka *et al*, 2015) or by TANK (Li *et al*, 2002), or the interaction between TNFR and TRAF6 inhibited by TRI4 (Moriya *et al*, 2015). Other inhibitors such as the TRAF2 inhibitor Liquidambaric acid (LDA), binds in the MATH/TRAF domain, preventing the binding of TRAF2 with β-Catenin (Yan *et al*, 2022), but the amino acids 408-431 of TRAF2 targeted by LDA are not homologous to those targeted on Traf4 by E(spl)m4. Other inhibitors could also interact outside of the MATH/TRAF and Coiled coil domains such as the interaction between TRAF3 and cIAP1 inhibited by the Staphylococcus aureus secreted extracellular fibrinogen-binding protein (Efb) (Zhang *et al*, 2022). Prediction with human TRAFs failed to detect any stable interaction between human TRAFs and the peptide identified in E(spl)m4, suggesting co-evolution constraints, but this new interaction could guide towards new strategies to design TRAF inhibitors by targeting the trimerization step.

Our studies suggest that E(spl)m4 can inhibit only the JNK activating role of Traf4 but not its role in the destabilization of AJ material. This latter role remain poorly understood, and might involve a potential E3 Ubiquitin Ligase activity of Traf4. Indeed, Traf4 has been shown to bind Armadillo (Mathew *et al*, 2011), and TRAF4, its human homologue has been found to bind to β-CATENIN and to promote its nuclear translocation and Wnt/β-Catenin pathway activation (Rozan & El-Deiry, 2006; Wang *et al*, 2014), thus potentially depleting the membrane-bound pool of β-Catenin. Alternatively, in mammals, both TRAF6 and TRAF3 can bind to c-SRC and other SFKs (Src Family Kinases), to control their Ubiquitination and activity (Wong *et al*, 1999; Liu *et al*, 2012; Johnsen *et al*, 2009). SFKs are important regulators of E-Cadherin-based AJs, exhibiting complex roles but mainly favoring AJs destabilization (reviewed in (Coopman & Djiane, 2016)). Whether overexpressed Traf4 could interact and/or control Src activity ultimately limiting the amount of E-Cad and Arm present at AJs, represents an attractive model, which remains to be tested. Since our modelling predicts that E(spl)m4 inhibits Traf4 by disrupting its trimerization, we propose that the two different activities of Traf4 depend on different assemblies of Traf4. First, Traf4 trimers formed upon interaction with clustered TNFR upstream receptor (or following overexpression), leads to JNK signaling. Second, Traf4 monomers resident of apical AJs control the abundance of AJ material, and in particular Arm/β-Cat. In this scenario, the binding of E(spl)m4 would have inhibitory effects only on the trimer and JNK pathway. While this model requires further validation, it is compatible with our observations.

E(spl)m4 represents thus an inhibitor for both Traf4 and the E3-Ubiquitin Ligase Neuralized. Interestingly, Traf4 and Neuralized have been implicated in the regulation of similar processes.

First, Neuralized controls the number of external sensory bristles (ESB) through its impact on Delta/Notch signaling and lateral inhibition (Lai *et al*, 2001; Yeh *et al*, 2001; Pavlopoulos *et al*, 2001; Le Borgne & Schweisguth, 2003). We report here that Traf4 controls the spacing between ESBs at the level of the wing margin (aka wing margin bristles), but we did not detect any alterations to Notch signaling. Strikingly, both Traf4 and its inhibitor E(spl)m4 are expressed in the proneural clusters which will give rise to ESBs (Lai *et al*, 2000; Reeves & Posakony, 2005; Preiss *et al*, 2001), and both can be regulated by Notch signaling. Indeed, we recently identified Traf4 as a direct Notch target in overgrowing neoplastic wing discs following constitutive Notch activation combined with epithelial polarity defects, with a Su(H) binding site located just upstream of the *Traf4-RA* promoter (Logeay *et al*, 2022). These observations suggest that, at least in the proneural clusters, complex cross-regulations between Notch, Traf4, E(spl)m4, and Neuralized take place.

Second, both Traf4 and Neuralized, have been implicated in AJ remodeling during embryonic mesoderm invagination (Mathew *et al*, 2009, 2011; Perez-Mockus *et al*, 2017; Chanet & Schweisguth, 2012). Thus the effects of Bearded proteins on mesoderm invagination and AJ remodeling (Chanet & Schweisguth, 2012; Perez-Mockus *et al*, 2017) might require re-interpretation taking into account a potential effect through Traf4. However, caution is required here since only E(spl)m4 and not all Bearded proteins has the potential to inhibit Traf4. Furthermore, while E(spl)m4 could prevent the JNK activation downstream of Traf4, it did not inhibit the effect of Traf4 on AJ proteins E-Cad and β-Catenin.

Further studies are thus required to better understand the relationships between Traf4, Neuralized, E(spl)m4, and the Notch pathway, and to explore whether peptides similar to E(spl)m4 could serve as natural inhibitors/modulators of other TRAF proteins, including in mammals.

## MATERIALS AND METHODS

### Drosophila genetics

All crosses were cultured at 25 or 29°C on standard food. UASt Traf4-PA:GFP was obtained by cloning *Traf4* amplified by PCR using the EST LD20987 as template, in frame with eGFP at the 3’ end.

New hypomorphic alleles for *Traf4* (ex40, ex65, ex111), were generated by imprecise excision of the P element EP578. Associated genomic deletions were mapped by PCR on genomic DNA (on homozygous individuals). The *Traf4^L2^* null allele was kindly gifted by Dr Yang (Wang *et al*, 2006). For cell competition experiments, *y-w-, hs-Flp, UAS CD8:GFP;; RpS3^Plac92^, Act>RpS3>Gal4/ TM6B* flies (kindly provided by Dr Eugenia Piddini) (Baumgartner *et al*, 2021), were crossed to flies carrying *UAS y-RNAi* controls or *UAS Traf4-RNAi* or *UAS E(spl)m4* transgenes. The progenies were heat shocked at 37C for 20 minutes at 48h after egg laying (ael) to allow for the excision of the FRT cassette and creating mosaic tissues with either one (loser, GFP positive) or two copies (winner, GFP negative) of the ribosomal protein coding gene *RpS3*. Wing discs were dissected at the wandering 3^rd^ instar larval stage (5days ael), and imaged to evaluate the amount of GFP-positive tissues.

Information on gene models and functions, and on *Drosophila* lines available were obtained from FlyBase (flybase.org – (Thurmond *et al*, 2019)). Stocks used are listed below:

Bloomington Drosophila Stock Center: BDSC
Vienna Drosophila Resource Center: VDRC

| Name / Gene | Transgene | Integration | Stock / Origin |
| --- | --- | --- | --- |
| **Gal4 Drivers** |  |  |  |
| *en-Gal4* | en2.4-Gal4[e16E] | on II | BDSC #30564 |
| *ptc-Gal4* | GawB ptc[559.1] | on II | BDSC #2017 |
| *sca-Gal4* | GawB sca[109-68] | on II | BDSC #6479 |
| **UAS Lines** |  |  |  |
| *Traf4* | Traf4[EP578] | on II | BDSC #17285 |
| *Traf4-RA:GFP* | Traf4-RA:GFP | on II | This study |
| *E(spl)m4* | E(spl)m4-BFM.A | on III | BDSC #26678 |
| *Tom* | Tom.L | on III | BDSC #26685 |
| *Traf4-RNAi* | KK107398 | VIE-260B | VDRC v110766 |
| *y-RNAi* | KK104196 | VIE-260B | VDRC v106068 |
| *w-RNAi* | HMS00045 | attP2 | BDSC #33644 |
| **Reporters** |  |  |  |
| *puc-LacZ* | puc[A251.1F3] | on III | BDSC #11173 |

### Yeast two hybrid

Yeast two-hybrid was used to test the interaction between Traf4-PA as bait and the different Brd proteins as preys. Traf4-PA and the different Brd genes were cloned from cDNA or genomic DNA for small intron-less ORFs, in the pAS2-1 (Gal4-BD, TRP1; Bait) or pACT2 (Gal4-AD, LEU2; Prey) vectors. Yeast cells were transfected with bait and prey plasmids using the Matchmaker kit (Clontech; according to manufacturer’s protocol). Transfected cells were plated on selective media lacking Leu and Trp for 3 days, before being streaked on media lacking Leu, Trp and Ade in order to test for potential interactions.

### Protein interaction modelling and molecular dynamics (MD) simulations

Structural models of Traf4 in complex with Bearded-family proteins were generated with the AlphaFoldC3 web server with default settings. The full-length proteins, as well as various fragments were tried as input sequences, and resulted in identification of the interaction between Traf4 MATH domain (residuesC328-480) and E(spl)m2 residuesC175-216 / E(spl)m4 residuesC100–136. For each pair five models were obtained; the model with the highest interface pLDDT was retained for analysis. To evaluate the potential impact of Traf4 self-association, an additional AlphaFoldC3 job was run with three identical Traf4 MATH +coiled coil domain sequences (trimeric input). Per residue pLDDT and predicted aligned-error (PAE) matrices were extracted from the JSON output. Interface residues were defined as any heavy atom within 4.5 Å of the partner chain in the top model. The two highest confidence complexes—Traf4-E(spl)m2(175-216) and Traf4-E(spl)m4(100-136)—were subjected to explicit-solvent MD in GROMACSC2024. Systems were described with the CHARMM27 all atom force field and solvated in a triclinic box (14 Å padding) filled with TIP3P water and 150 mM NaCl. Energy minimization (steepest descent, <1000CkJCmol⁻¹Cnm⁻¹) was followed by 500Cps of equilibration at 300 K (v-rescale) and 1Cbar (Parrinello–Rahman barostat). Production runs comprised one 1 µs trajectory per complex. Interface RMSD (iRMSD) was calculated after backbone alignment of Traf4 residuesC328–480 using *gmx rms*. BSA over time was obtained with *gmx sasa*.

### Immunofluorescence

3^rd^ instar wing imaginal discs were dissected in PBS, fixed in 4% formaldehyde for 20min (discs) at room temperature (RT), washed 3x 10min in PBT (PBS 0,2% Triton X-100), and blocked in PBT-BSA (PBT 0.5% BSA) for 30min. Primary antibodies were incubated overnight at 4°C in PBT-BSA. Tissues were then washed 3x 10min in PBT, and secondary antibodies were incubated 90min at RT in PBT-BSA. They were washed 3×10 min in PBT and mounted in CitiFluor^TM^ AF1 (Agar). Images were acquired on a Leica SP5 confocal microscope or on an upright Leica THUNDER or a Zeiss Apotome microscope.

Primary antibodies were: Mouse anti-Armadillo (1:25; Developmental Study Hybridoma Bank – DSHB #N2 7A1), Rabbit anti-cleaved Caspase 3 (D175) (Cell Signaling Technology – CST #9661; 1/500), Rabbit anti-Cleaved Dcp-1 (1:500; CST #9578), Mouse anti-Discs large (1:25; DSHB #4F3), Rat anti-E-Cadherin (1:25; DSHB #DCAD2), Mouse anti-b-Galactosidase (1:25; DSHB #40-1a), Rabbit anti-GFP (1:500; Torrey Pines Biolabs #TP401), Rabbit anti-aPKC (anti-PKCz C-20, SantaCruz; 1:500). Secondary antibodies conjugated to Alexa Fluor 488, 555, 647, or to Cy3 were from Jackson Labs Immuno Research (1:200).

Images were acquired on a Leica 510 confocal microscope or on a Leica Thunder epifluorescence microscope. Images were then treated using FIJI.

## ACKNOWLEDGEMENTS

We thank M. Leptin, E. Piddini, and X. Yang for sharing flies and reagents. We acknowledge the Bloomington Drosophila Stock Center (BDSC - NIH P40OD018537), the Vienna Drosophila Resource Center (VDRC), the Developmental Studies Hybridoma Bank, the Montpellier Drosophila facility, the Montpellier Imaging facility (MRI), and FlyBase for providing reagents and tools critical for our research.

## AUTHOR CONTRIBUTIONS

Conceptualization: AD and SJB. Methodology: PL, CL, AD. Validation: PL, CL, RL, KS, AD. Formal analysis: CL, AD. Investigation: PL, CL, RL, KS, AD. Writing – original Draft: AD. Writing – review and editing: PL, SJB, AD. Visualization: PL, CL, AD. Supervision: AD. Project administration: AD. Funding acquisition: SJB, AD.

## DATA AVAILABILITY

No new dataset was generated for this study.

---

**Supplemental Figure S1. Overexpressed Traf4 and apico-basal polarity markers (related to Fig. 1)**

**A-B.** Staining of 3^rd^ instar wing imaginal disc with overexpressed Traf4-PA:GFP in the posterior compartment showing the Traf4-PA:GFP overexpression domain (GFP, green in A&B), the subapical compartment marker aPKC (red in A, white in A’), the Septate Junctions marker Discs Large (Dlg, red in B, white in B’) and E-Cadherin (E-Cad, blue in A&B, white in A’’&B’’).

**Supplemental Figure S2. Interaction between Traf4 and E(spl)m2 and E(spl)m4 (related to Fig. 4)**

**A.** Alignment of the different Bearded proteins using the Uniprot portal and highlighting similarity. The red boxes represent previously identified motifs, where motif 1 corresponds to an amphipathic alpha helix, and motif 2 is the domain mediating the interaction between Brd proteins (except E(spl)m2) and the E3-Ubiquitin Ligase Neuralized. This study identifies the motif 3 as the interface with Traf4.

**B.** Comparison of Traf4-E(spl)m4 predicted interaction interface with interfaces observed on other TRAF family members complexed with various proteins and peptides. Pdb codes 1QSC, 1CA9, 1F3V, 4GHU and 1KZZ were used.

**Supplemental Figure S3. E(spl)m4 suppresses Traf4-induced cell delamination (related to Fig. 5)**

**A-D.** Staining of 3^rd^ instar wing imaginal disc overexpressing Traf4-RA:GFP at 29°C either with GFP (A&B) or with E(spl)m4 (C&D) and showing the overexpression domain (GFP, green in A-D, white in A’-D’), E-Cadherin (blue in A-D), and cleaved Caspase 3 (Cas3, red in A-D, white in A’’-D’’). Discs are shown along the xy axis in A&C and along the z axis in B&D.

