## Supplementary figures and images for "E(spl)m4 Directly Antagonizes Traf4 to Inhibit JNK Signaling in *Drosophila*"

### Supplemental Fig. S1

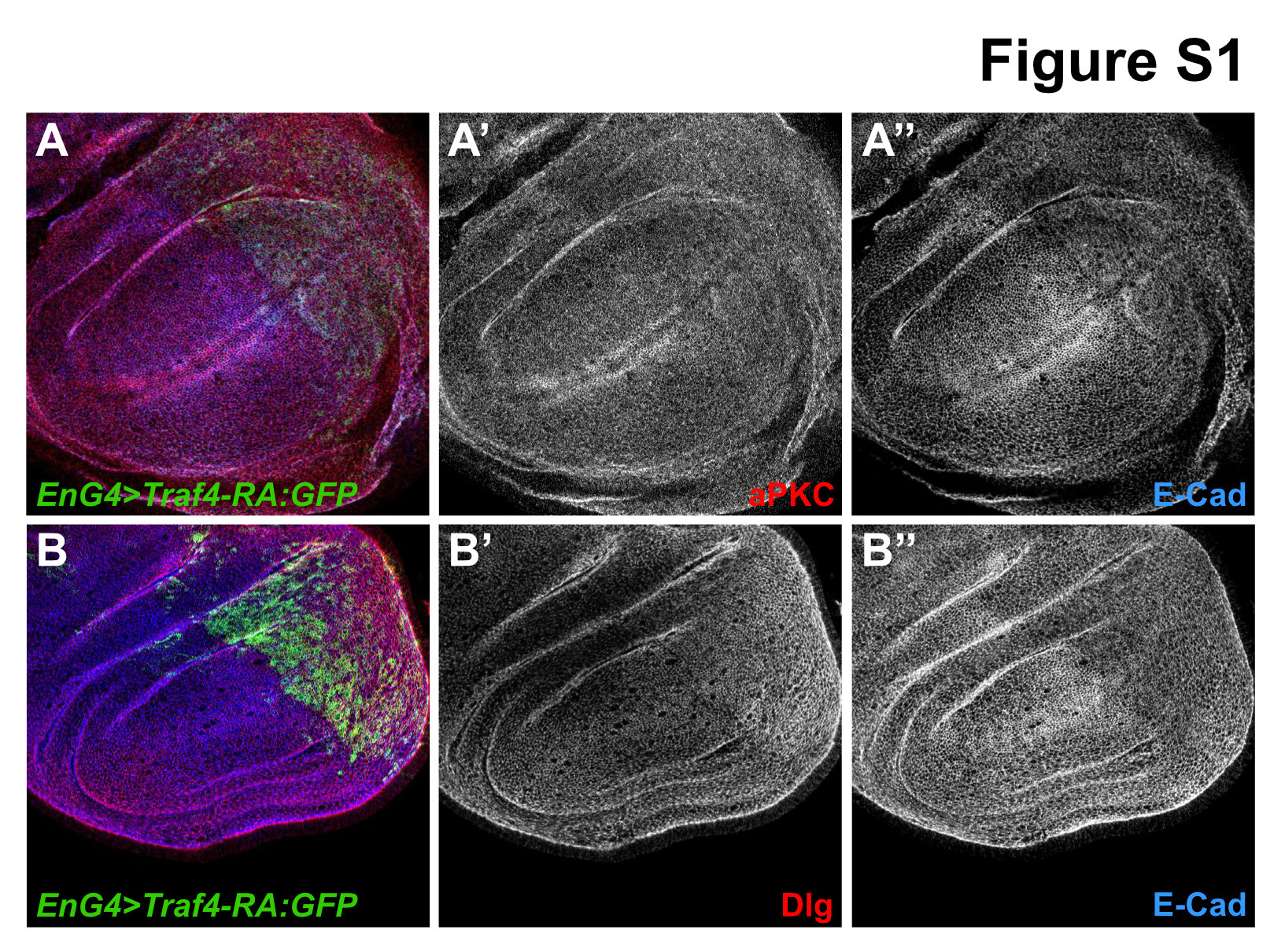

### Supplemental Fig. S2

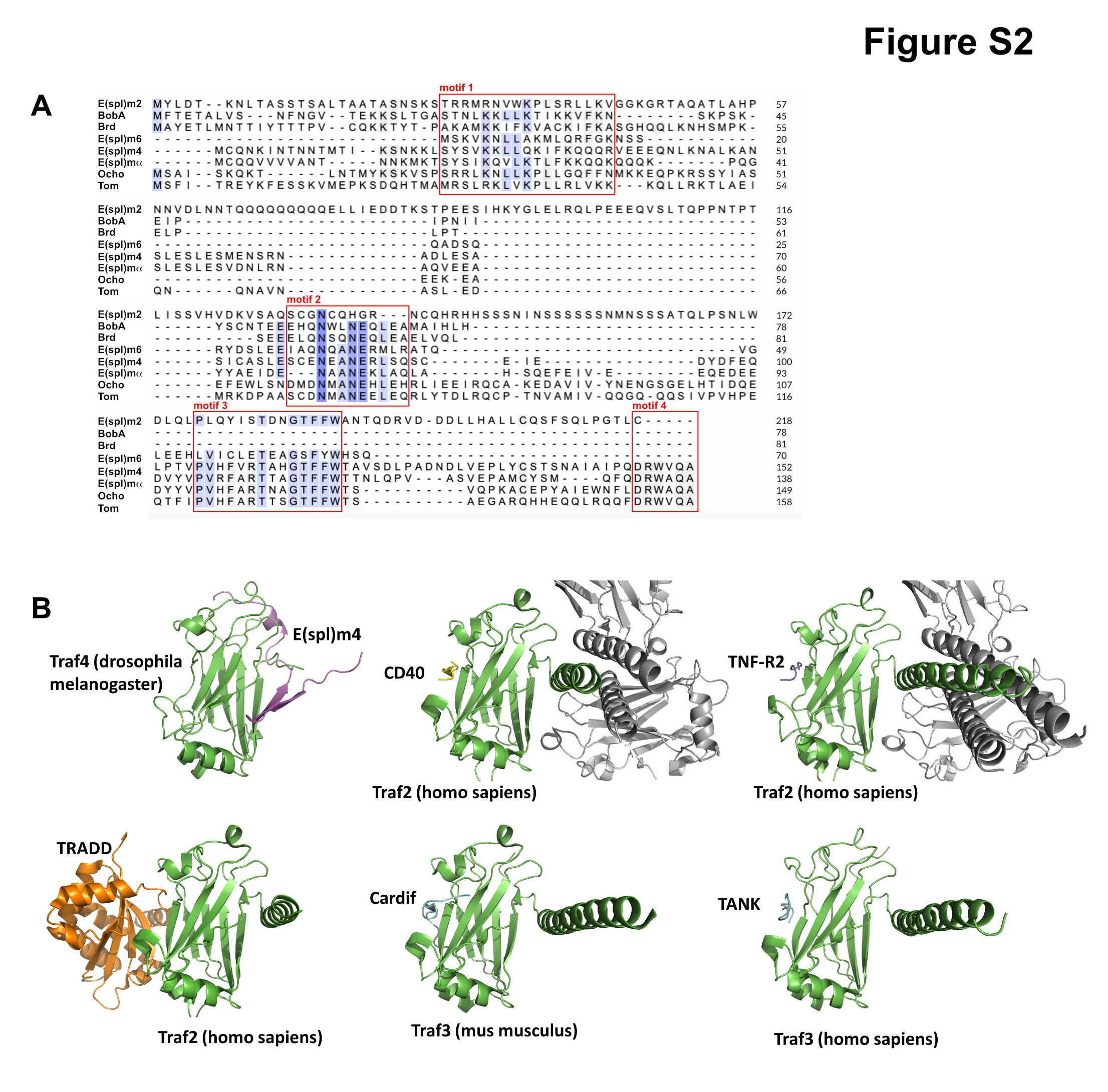

### Supplemental Fig. S3

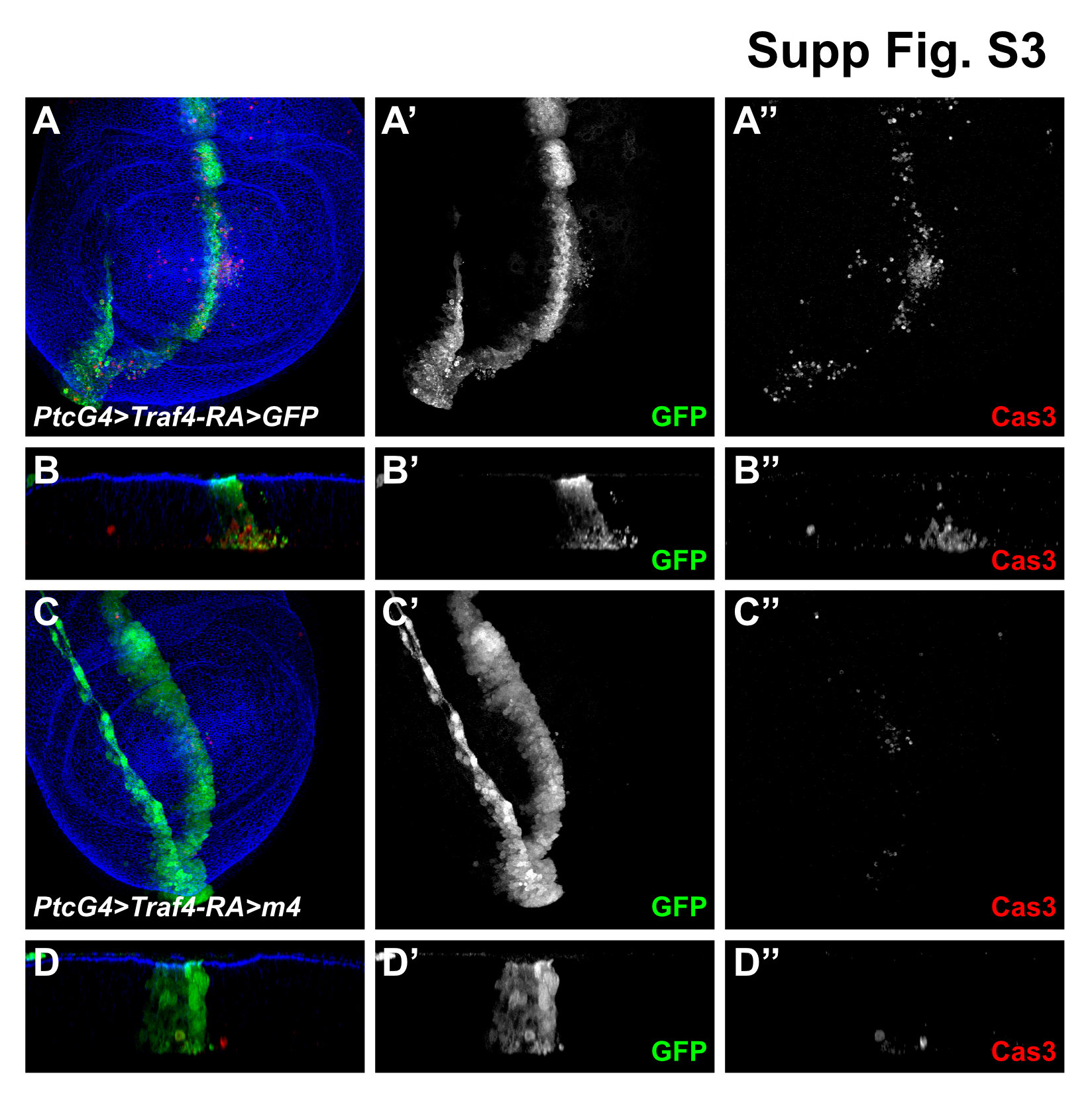
